# Identifying miRNA synergism using multiple-intervention causal inference

**DOI:** 10.1101/652180

**Authors:** Junpeng Zhang, Vu Viet Hoang Pham, Lin Liu, Taosheng Xu, Buu Truong, Jiuyong Li, Nini Rao, Thuc Duy Le

## Abstract

**Background:** Studying multiple microRNAs (miRNAs) synergism in gene regulation could help to understand the regulatory mechanisms of complicated human diseases caused by miRNAs. Several existing methods have been presented to infer miRNA synergism. Most of the current methods assume that miRNAs with shared targets at the sequence level are working synergistically. However, it is unclear if miRNAs with shared targets are working in concert to regulate the targets or they individually regulate the targets at different time points or different biological processes. A standard method to test the synergistic activities is to knock-down multiple miRNAs at the same time and measure the changes in the target genes. However, this approach may not be practical as we would have too many sets of miRNAs to test.

**Results:** In this paper, we present a novel framework called miRsyn for inferring miRNA synergism by using a causal inference method that mimics the multiple-intervention experiments, e.g. knocking-down multiple miRNAs, with observational data. Our results show that several miRNA-miRNA pairs that have shared targets at the sequence level are not working synergistically at the expression level. Moreover, the identified miRNA synergistic network is small-world and biologically meaningful, and a number of miRNA synergistic modules are significantly enriched in breast cancer. Our further analyses also reveal that most of synergistic miRNA-miRNA pairs show the same expression patterns. The comparison results indicate that the proposed multiple-intervention causal inference method performs better than the single-intervention causal inference method in identifying miRNA synergistic network.

**Conclusions:** Taken together, the results imply that miRsyn is a promising framework for identifying miRNA synergism, and it could enhance the understanding of miRNA synergism in breast cancer.

## Background

MicroRNAs (miRNAs) are a class of short non-coding RNAs with ~23 nucleotides (nts) in length. They play an important regulatory role at the post-transcriptional level by targeting messenger RNAs (mRNAs) for degradation or translation repression [1]. Previous studies have demonstrated that miRNAs play a crucial role in regulating gene expression involved in diverse biological processes, including cell proliferation, cell death, cell apoptosis and human cancers [2–4]. Generally, the relationships between miRNAs and their target genes are not one-to-one but many-to-many, indicating cooperative regulation of miRNAs. The co-regulation of miRNAs has been accepted and confirmed by cross-linking and immunoprecipitation technologies, and may be related to human complex diseases [5]. Therefore, studying miRNA synergism can greatly help to understand the synergistic regulation mechanism of miRNAs in human diseases.

Until now, a number of computational methods have been proposed to identify miRNA synergism. These methods can be divided into three different categories: (1) sequence-based [6–8], (2) correlation-based [9–14], and (3) causality-based [15]. In the first category, the sequence data mainly includes putative miRNA-target interactions and protein-protein interactions (PPIs). For each candidate miRNA-miRNA pair, the methods firstly evaluate the significance of common target genes by using a hypergeometric test. Then, by conducting Gene Ontology (GO) [16] or Kyoto Encyclopedia of Genes and Genomes (KEGG) [17] enrichment analysis of the shared target genes, they determine whether a candidate miRNA-miRNA pair is functionally synergistic or not. The main limitation of the methods in this category is that they only use static data in the study of miRNA synergism. In fact, the co-regulation between miRNAs is usually dynamic in human cancers [18]. Methods in the second category use expression data of miRNAs to identify differentially expressed miRNA synergistic network or integrate matched miRNA and mRNA expression data with sequence data to infer miRNA synergistic networks. However, the identified miRNA synergistic networks or modules using statistic correlation methods may not reveal the causal relationships of gene regulation. To address this issue, a causality-based method [15] (the third category) has been presented to infer miRNA-target causal relationships. The method only simulates the causal effect in single-intervention experiments, e.g. knocking-down a single miRNA each time. However, miRNA co-regulation simultaneously involves multiple miRNAs.

In general, the miRNA-miRNA synergistic pairs identified by several existing methods at the sequence level may not crosstalk with each other to co-regulate target genes at the expression level. Previous study [19] has shown that miRNAs tend to synergistically control expression levels of target genes. It is necessary to integrate expression data for identifying miRNA-miRNA synergistic pairs at the expression level. Moreover, all existing approaches don’t explicitly look at “simultaneous” co-regulation of multiple miRNAs on the target genes, e.g. causal effects of multiple synergistic miRNAs on the shared target genes.

To address the above issues, in this work, we present a framework called miRsyn for inferring <u>miR</u>NA <u>syn</u>ergism from both sequence-based binding information and expression data by simulating multiple-intervention experiments, e.g. knocking-down multiple miRNAs at the same time. We apply the proposed method to The Cancer Genome Atlas (TCGA) breast cancer dataset. The results show that several miRNAs that have shared targets at the sequence level may not be working synergistically at the expression level, and the miRNA synergistic modules discovered are strongly related to breast cancer. Our further analyses also reveal that most of synergistic miRNA-miRNA pairs tend to be co-expressed, which help make a rapid response to external disturbances. Finally, the comparison results demonstrate that multiple-intervention causal inference method performs better than single-intervention causal inference method in studying miRNA synergism.

## Methods

### Overview of miRsyn

As illustrated in Figure 1, miRsyn is a step-wise method for identifying miRNA synergism. Firstly, given matched miRNA and mRNA expression data, we use feature selection based on the Cox regression model [20] to identify significant miRNAs and mRNAs. Then, by using multiple-intervention causal inference method [21], we obtain joint causal effects between the significant miRNAs and mRNAs. At the same time, the putative miRNA-target binding information is used to generate regulatory relationships between significant miRNAs and mRNAs. By integrating joint causal effects and binary relationships between significant miRNAs and mRNAs, we find a set of miRNAs with the maximum joint causal effect on each mRNA. The miRNAs in each set of miRNAs synergistically regulate their target mRNAs, and all synergistic miRNA-miRNA pairs are combined to construct the miRNA synergistic network. To identify miRNA synergistic modules, we firstly find a set of bi-cliques with at least 2 miRNAs and mRNAs based on putative miRNA-mRNA binding information. For each bi-clique, the subset of bi-clique with the maximum joint causal effect is regarded as a miRNA synergistic module. Finally, we conduct functional analysis of miRNA synergism at both network and module levels.

In the following, we will describe the key steps in detail.

**Figure 1.**
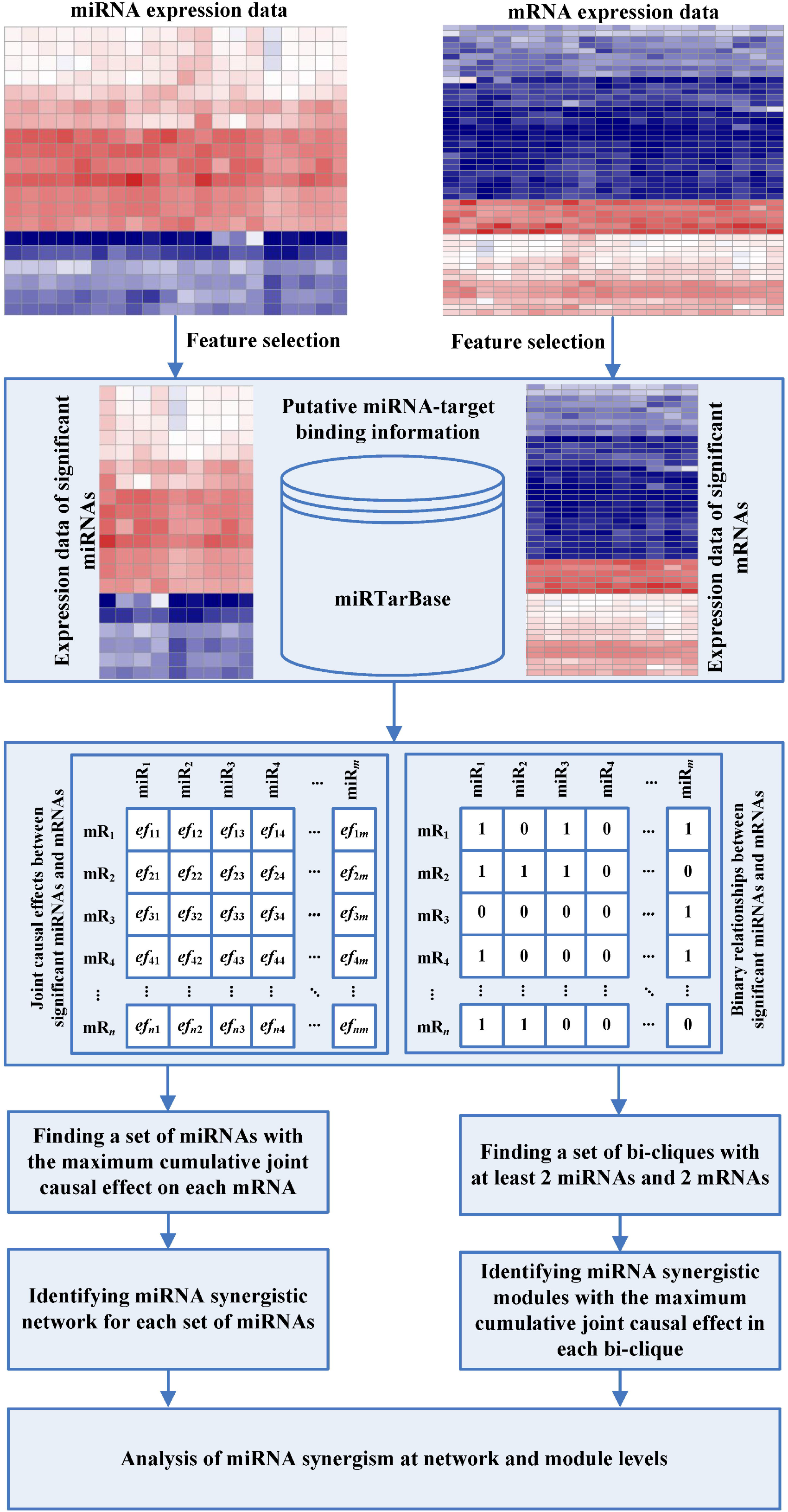
The workflow of miRsyn. The process contains three main steps. Firstly, we identify significant miRNAs and mRNAs using feature selection from miRNA and mRNA expression data. Secondly, by integrating expression data of significant miRNAs and mRNAs and putative miRNA-target interactions, we identify miRNA synergistic network and modules. Finally, we make a functional analysis of the identified miRNA synergistic network and modules.

### Estimating multiple-intervention effects

Let *G* = (*V*, *E*) be a graph including a set of vertices *V* and a set of edges *E* ⊆ *V* × *V* . Here, *V*={*X*_1_, …, *X*_*p*_, *Y*_1_, …,*Y*_*q*_} is a set of random variables denoting the expression levels of *p* miRNAs and *q* mRNAs, and *E* represents the regulatory relationships between these variables. If a graph *G* only contains directed edges and no cycles, the graph is a Directed Acyclic Graph (DAG). In DAG *G*, if there is an edge *X*_*i*_ → *Y*_*j*_, *X*_*i*_ (*i* ∈ {1,…, *p*}) is a parent of *Y*_*j*_ (*j* ∈ {1,…, *q*}) and *Y*_*j*_ is a child of *X*_*i*_◻. The DAG *G* is a causal DAG if and only if *X*_*i*_ is a direct cause of *Y*_*j*_ [22]. Following the Markov Assumption that a node in a DAG is conditionally independent of its non-descendants, given its parents, a DAG encodes a distribution, as a product of the conditional probabilities of all nodes. A DAG can be read out as a set of conditional independent relationships of variables. An *equivalence class* of DAGs, which encodes the same conditional independencies in a given data, can be described by a *completed partially directed acyclic graph* (CPDAG) which includes both directed edges and undirected edges [23].

Let’s assume that we have observational data (e.g. gene expression data) that are multivariate Gaussian and faithful to the true (but unknown) underlying causal DAG without hidden variables. Under the assumption, the Joint-IDA (Joint Intervention calculus when the DAG is Absent) [21] estimates the multiset of possible total joint effect of *X* ({*X*_1_,…, *X*_*p*_}) on *Y*_*j*_ (*j* ∈ {1,…, *q*}). The total effect of *X* on *Y*_*j*_ in a joint intervention on *X*_*k*_ (*k* ≠ *i*) is denoted by (*ef*_1*j*_, *ef*_2*j*_, …, *ef*_*pj*_), where *ef*_*ij*_ represents the direct causal effect of *X*_*i*_ (*i* ∈ {1,…, *p*}) on *Y*_*j*_, when keeping intervention values of other variables *X*_*k*_ constant. The joint effect (*ef*) of *X* on each of *Y*_*j*_ is formally defined as follows:

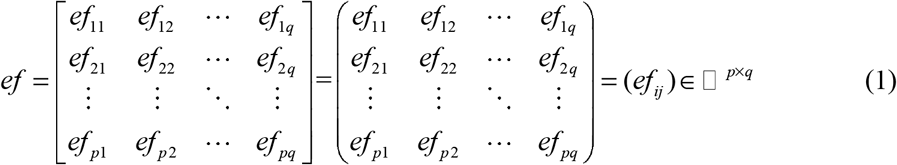

Where

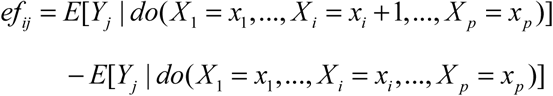

In the formula, *do*(.) is the ‘do’ operation to set *X_i_* to a value, e.g. (*x*_*i*_ + 1) or *x*_*i*_ (*i* ∈{1,…, *p*}), and this mimics a real world manipulation by setting a variable to a value *x*_*i*_. *E*[.] is the expectation of variable *Y*_*j*_ when variable *x*_*i*_ is manipulated and other variables *X*_*k*_ (*k* ≠ *i*) are held constant.

Joint-IDA implemented in the R package *pcalg* [24] can be directly used to calculate joint casual effect, but is not applicable to gene expression datasets with thousands of variables. We have implemented a parallelized Joint-IDA algorithm in the R package, *ParallelPC* [25] which uses a multiple-core CPU to speed up the runtime of the Joint-IDA algorithm.

Let us consider a subset of miRNAs ({*X*_1_, …, *X*_*m*_}) where *m* ≤ *p* . We are interested in the cumulative joint causal effect of the *m* miRNAs on mRNA *Y*_*j*_ when the *m* miRNAs are knocked off where other miRNAs are kept the intervention values constant. The cumulative joint causal effect(δ_*j*_) of *m* miRNAs on each of *Y*_*j*_ is defined in the following:

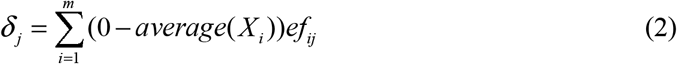

where *ef*_*ij*_ represents the amount of *Y*_*j*_ change due to a unit change of *X*_*i*_, *average*(*X*_*i*_) denotes the average expression level of *X*_*i*_ in the expression data, and (0 - *average*(*X*_*i*_)) indicates ‘knocking off’ miRNA *X*_*i*_ completely.

A DAG cannot be uniquely identified from data, and instead, an equivalence class of DAGs is identified. We estimate a multiset of possible cumulative joint causal effects using the set of equivalent DAGs. The maximum of cumulative joint causal effects in the multiset is reported as the estimated cumulative joint causal effect.

### Identifying miRNA synergistic network

After feature selection in the matched miRNA and mRNA expression data, we obtain a list of significant *p* miRNAs and *q* mRNAs. Given the putative miRNA-target binding information, let *A*={*miR*_1_,…, *miR*_*p*_} be a set of significant miRNAs that have binding sites with a significant target mRNA *mR*_*j*_ (*j* ∈ {1,…, *q*}). Our aim is to find a set of miRNAs 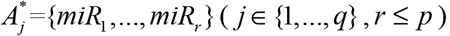, which has the maximum cumulative joint causal effect on mRNA *mR*_*j*_. This is the estimated cumulative joint causal effect of 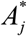 on the mRNA *mR*_*j*_ when all miRNAs in 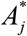 are knocked off at the same time in data. In each set of 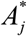, the miRNAs synergistically regulate *mR*_*j*_, and form a miRNA-miRNA synergistic sub-network. All miRNA-miRNA sub-networks are then integrated to maximal miRNA synergistic networks. Our identified miRNA synergistic networks are different from those obtained from existing methods, as we draw each 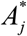 as a sub-network by simulating multiple gene knock-down experiments.

### Identifying miRNA synergistic modules

We firstly initialize the miRNA-mRNA bipartite network between the *p* significant miRNAs and *q* significant mRNAs by using putative miRNA-target binding information. Then, we use the R package *biclique* [26] to find all the bi-cliques from the miRNA-mRNA bipartite network. The bi-cliques provide the candidate miRNA synergistic modules for testing the miRNA synergistic activities. For a bi-clique, let *C*={*miR*_1_,…, *miR*_*r*_} (subset of *p* significant miRNAs) and *D*={*mR*_1_,…, *mR*_*l*_}(subset of *p* significant mRNAs) denote *r* (≥ 2) miRNAs and *l* (≥ 2) mRNAs in the bi-clique. Based on the joint causal effects of *C* on each mRNA of *D*, we find a set of *C\** ={*miR*_1_, …, *miR*_*l*_}(subset of *C*) and *D\**={*mR*_1_, …, *mR*_l_} (subset of *D*) with the maximum cumulative joint causal effect between *C*\* on every mRNA in *D*\*. The identified (*C*\*, *D*\*) is regarded as a miRNA synergistic module where the number of miRNAs or mRNAs is at least 2.

### Topological and functional analysis of miRNA synergism

The topological analysis of miRNA synergism could help to understand the internal organization of miRNA synergistic network, e.g. power law degree distribution, the average clustering coefficient and the average characteristic path length. If the node degree in a biological network obeys a power law curve (in the form of *y* = *bx*^*a*^) distribution with high value of *R*^2^, the network is regarded to be scale-free. Here, the *R*^2^ value is a deterministic coefficient to measure the quality of a power curve fit. The interval of *R*^2^ value is [0 1]. A larger *R*^2^ value indicates a better power law curve fit. The average clustering coefficient is used to evaluate the dense neighborhood of a biological network. In a small-world biological network, the average characteristic path length is much larger than that of random networks [27, 28]. The average characteristic path length indicates the density of a biological network. In a small-world biological network, the average characteristic path length is smaller than that of random networks [28].

In this work, we obtain topological features (power law degree distribution, the average clustering coefficient and the average characteristic path length) of the miRNA synergistic network by using the *NetworkAnalyzer* plugin [29] in Cytoscape [30]. For generating random networks, we use the duplication model [31] of the *RandomNetworks* plugin (https://github.com/svn2github/cytoscape/tree/master/csplugins/trunk/soc/pjmcswee/src/cytoscape/randomnetwork) in Cytoscape. We construct 10000 random instances by randomizing the miRNA synergistic network, and calculate the average clustering coefficient and the average characteristic path length of networks.

We conduct functional enrichment analysis to investigate the biological functions of miRNA synergism. For the identified miRNA synergistic network, we use the online tool miEAA [32] to infer the significantly enriched or depleted biological processes, pathways and diseases associated with synergistic miRNAs (*p*-value < 0.05). For the identified miRNA synergistic modules, we focus on annotating breast cancer related miRNA synergistic modules by conducting breast cancer enrichment analysis. Here, we use a hypergeometric test to perform breast cancer enrichment analysis. For each miRNA synergistic module, the significance *p*-value of breast cancer genes is calculated as follows.

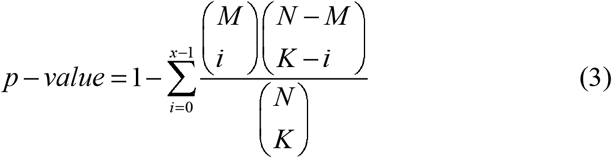

In the formula, *N* is the number of significant genes (including miRNAs and mRNAs) after feature selection, *M* denotes the number of breast cancer genes in significant genes, *K* represents the number of genes in each miRNA synergistic module, and *x* is the number of breast cancer genes in each miRNA synergistic module. The miRNA synergistic modules with *p*-value < 0.05 are regarded as breast cancer related modules.

## Results

### Data source

We obtain the matched breast cancer expression data of miRNAs and mRNAs and the clinical information of breast cancer samples from TCGA [33]. Firstly, we remove all the male samples for breast cancer because this is a relative minority event. For the matched miRNA and mRNA expression data, the genes with missing values across the samples (>30%) is removed. The remaining missing values are imputed using the *k*-nearest neighbours (KNN) algorithm from the *impute* R package [34]. Then, we conduct *log*_2_ (*x* +1) transformation and *z*-score normalization for the expression levels of miRNAs and mRNAs. In addition, we use the *miRBaseConverter* R package [35] to convert miRNA names to the latest version of miRBase. Finally, we use the *FSbyCox* function (a feature selection based on Cox regression model) from the *CancerSubtypes* R package [36] to identify significant miRNAs and mRNAs. After the feature selection, we identify expression data of 79 miRNAs and 1314 mRNAs in 753 breast cancer samples at a significant level (*p*-value < 0.05) in total. For the putative miRNA-target interactions, we use the experimentally validated interactions from miRTarBase v7.0 [37]. A list of breast cancer related miRNAs are obtained from HMDD v3.0 [38], miR2Disease [39], miRCancer [40] and oncomiRDB [41]. A list of breast cancer related genes is obtained from DisGeNET v5.0 [42] and COSMIC v86 [43].

### MiRNA synergistic network is small-world and biologically meaningful

By following the steps of Figure 1, we have identified a list of 702 miRNA-miRNA synergistic pairs between 78 miRNAs (details can be seen in Additional file 1). These miRNA-miRNA synergistic pairs are integrated into a miRNA synergistic network. Out of the 78 miRNAs, the number of breast cancer related miRNAs is 39 (red nodes in Figure 2). The hub miRNAs with higher degrees in miRNA synergistic network tend to be essential. In this work, 8 miRNAs with higher degrees (about 10% of miRNAs in miRNA synergistic network) are regarded as hub miRNAs. Except one hub miRNA (miR-186-5p), 7 hub miRNAs (miR-10a-5p, and miR-150-5p, miR-192-5p, miR-26a-5p, miR-301a-3p, miR-484, miR-98-5p) are breast cancer related miRNAs. This result indicates that most of hub miRNAs are breast cancer causal miRNAs. We define that breast cancer related miRNA-miRNA pairs are those in which the two synergistic parties are breast cancer related miRNAs. As a result, we obtain a list of 269 breast cancer related miRNA-miRNA pairs (details can be seen in Additional file 1).

**Figure 2.**
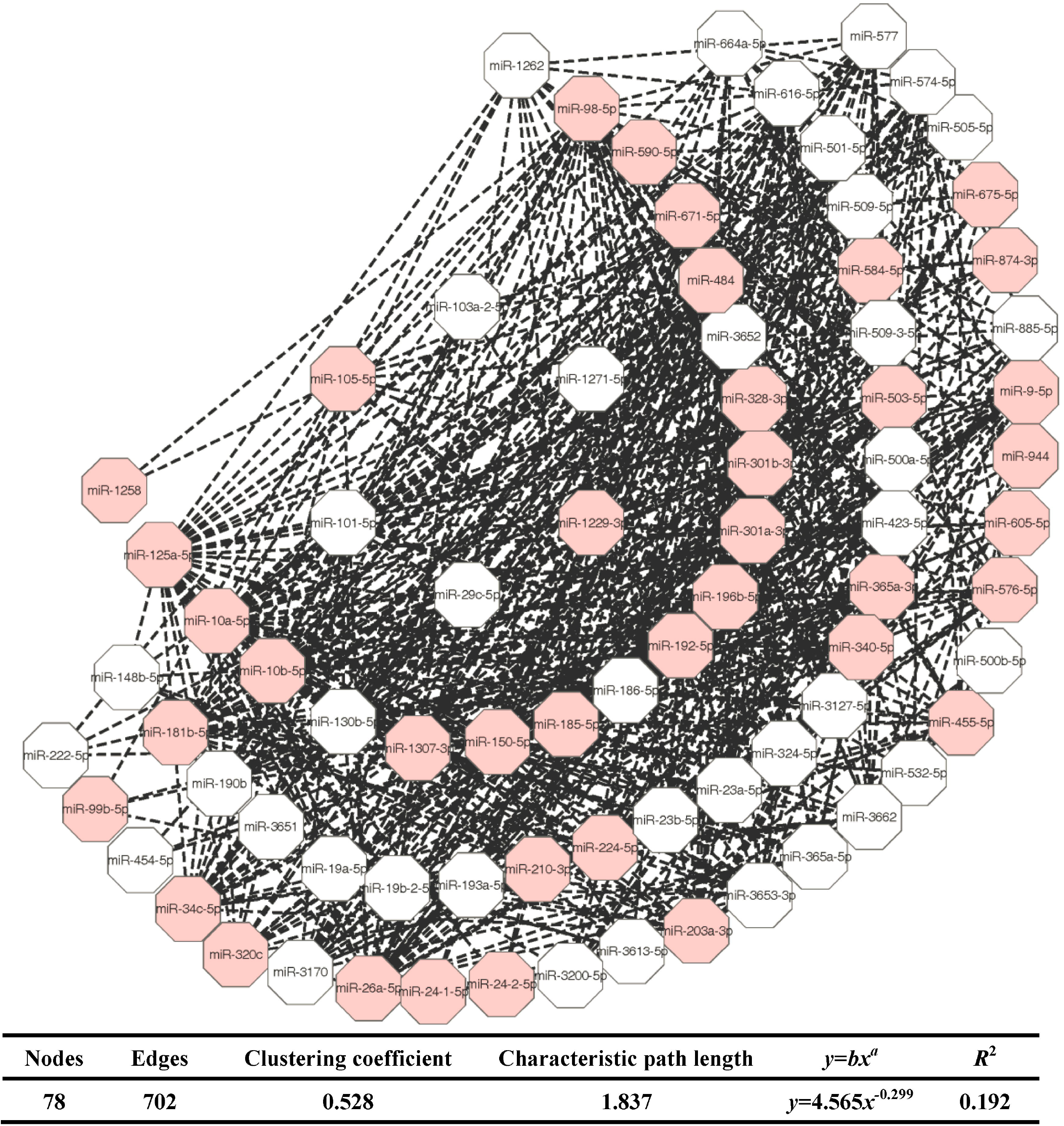
Visualization of miRNA synergistic network generated by Cytoscape. The breast cancer related miRNA nodes are colored in red, and the non breast cancer related miRNA nodes are colored in white. The dash lines denote synergistic relationships.

As shown in Figure 2 (table at the bottom of the figure), the distribution of node degrees of the miRNA synergistic network do not follow power law distribution with *R*^2^ = 0.192. This result indicates that the identified miRNA synergistic network is not scale-free. However, the miRNA synergistic network exhibits dense local neighborhoods with the average clustering coefficient of 0.528, which is much larger than that of random networks (0.178 ± 0.037). In addition, the miRNAs in the network are closely connected with the average characteristic path length of 1.837, which is smaller than that of random networks (2.511 ± 0.048). Altogether, the dense local neighborhoods and the small average characteristic path length imply that the miRNA synergistic network is small-world, and can be used to predict miRNA synergism [27, 28].

To investigate the potential biological processes, pathways and diseases related to the synergistic miRNAs, we conduct functional enrichment analysis of them. As shown in Table 1, the synergistic miRNAs are significantly enriched in several biological processes, pathways and diseases associated with breast cancer, such as cell cycle (GO0007050, GO0007093) [44],cell division (GO0051781) [45], cell apoptosis (GO0002903, GO0042981, GO0043065, hsa04210) [46], cell migration (GO0030334, GO0010595, GO0030335,) [47], cell differentiation (GO0045595, GO0045446,), cell proliferation (GO0050678, GO0072091) [49], signaling pathway (P00038, P00056, WP304) [50] and Breast Neoplasms. The detail information of miRNA enrichment analysis results can be seen in Additional file 2. This result demonstrates that the miRNA synergistic network is closely associated with the biological condition of breast cancer dataset, and is biologically meaningful.

**Table 1.** A portion of enriched or depleted biological processes, pathways and diseases associated with breast cancer by using miRNA enrichment analysis.

| Category | Subcategory | Enrichment | *p*-value | #miRNAs |
| --- | --- | --- | --- | --- |
| Gene Ontology | GO0007050:cell cycle arrest | enriched | 3.079E-02 | 27 |
|  | GO0007093:mitotic cell cycle checkpoint | enriched | 8.732E-03 | 10 |
|  | GO0051781:positive regulation of cell division | enriched | 1.146E-02 | 16 |
|  | GO0002903:negative regulation of b cell apoptotic process | depleted | 3.023E-02 | 3 |
|  | GO0042981:regulation of apoptotic process | enriched | 2.025E-02 | 21 |
|  | GO0043065:positive regulation of apoptotic process | enriched | 7.562E-03 | 27 |
|  | GO0030334:regulation of cell migration | enriched | 3.296E-02 | 14 |
|  | GO0010595:positive regulation of endothelial cell migration | enriched | 2.851E-02 | 12 |
|  | GO0030335:positive regulation of cell migration | enriched | 2.327E-02 | 20 |
|  | GO0045595:regulation of cell differentiation | enriched | 3.939E-02 | 10 |
|  | GO0045446:endothelial cell differentiation | depleted | 3.330E-04 | 2 |
|  | GO0050678:regulation of epithelial cell proliferation | enriched | 3.845E-02 | 3 |
|  | GO0072091:regulation of stem cell proliferation | enriched | 3.497E-02 | 2 |
|  | GO0010719:negative regulation of epithelial to mesenchymal transition | depleted | 4.996E-02 | 3 |
| Pathways | hsa04210:Apoptosis | enriched | 3.859E-02 | 22 |
|  | P00038:JAK STAT signaling pathway | enriched | 4.995E-03 | 2 |
|  | P00056:VEGF signaling pathway | enriched | 3.376E-02 | 15 |
|  | WP304:Kit receptor signaling pathway | enriched | 7.517E-03 | 18 |
| Diseases | Breast Neoplasms | enriched | 7.184E-03 | 11 |

### A number of miRNA synergistic modules are significantly enriched in breast cancer

We have identified 361 miRNA synergistic modules by following the steps in Figure 1 (details in Additional file 3). To understand whether the identified miRNA synergistic modules are closely associated with breast cancer, we conduct breast cancer enrichment analysis of these modules. As a result, the number of miRNA synergistic modules significantly enriched in breast cancer is 72 (*p*-value < 0.05), indicating that a number of miRNA synergistic modules is closely related to the biological condition of breast cancer dataset (details in Additional file 3).

### Most of synergistic miRNA-miRNA pairs show the same expression patterns

In this study, we use Pearson correlation of each synergistic miRNA-miRNA pair to measure the co-expression level. A synergistic miRNA-miRNA pair with significantly positive correlation (*p*-value < 0.05) is regarded as a co-expressed pair. Out of 702 synergistic miRNA-miRNA pairs, we discover that 499 synergistic miRNA-miRNA pairs are co-expressed (details in Additional file 4). This result indicates that most of synergistic miRNA-miRNA pairs (~71.08%) show similar expression patterns. It also implies that most miRNAs with similar expression patterns would like to collaborate with each other to co-regulate target genes. The result is consistent with previous studies [7, 51].

### Several synergistic miRNA-mRNA pairs at the sequence level are not working synergistically at the expression level

At the sequence level, we only use putative miRNA-target interactions to construct miRNA synergistic network. In this work, we use the DmirSRN motif in [15] to generate miRNA synergistic regulatory network. Consequently, we find that 1313 miRNA-miRNA pairs can directly regulate the same target by cooperating with each other at the sequence level (details in Additional file 5). Out of 1313 synergistic miRNA-miRNA pairs at the sequence level, 611 miRNA-miRNA pairs are not working synergistically at the expression level by comparing with the miRNA synergistic network generated by miRsyn (details in Additional file 5). This result implies that miRNA-miRNA pairs that have shared targets at the sequence level may not work synergistically at the expression level.

### Comparison results

There are several existing methods to infer miRNA synergistic network by using different types of datasets. However, to have a fair comparison (i.e. using the same data types and similar inference method for estimating causal effects of miRNAs on mRNAs), we focus the comparison on one existing method mirSRN [15] only.

The comparison result of our method miRsyn with mirSRN is shown in Figure 3. The detailed results of mirSRN can be seen in Additional file 6. In terms of the identified miRNA synergistic pairs (Figure 3A), the number of miRNA synergistic pairs predicted by miRsyn (702) is more than that by mirSRN (239). The majority of the identified miRNA synergistic pairs by mirSRN (163) are predicted by miRsyn. As for the significantly enriched terms (Gene Ontology, Pathways and Diseases) associated with the identified miRNA synergistic network (Figure 3B), the identified miRNA synergistic network from miRsyn is significantly enriched in more number of functional terms excepting the terms of Diseases.

**Figure 3.**
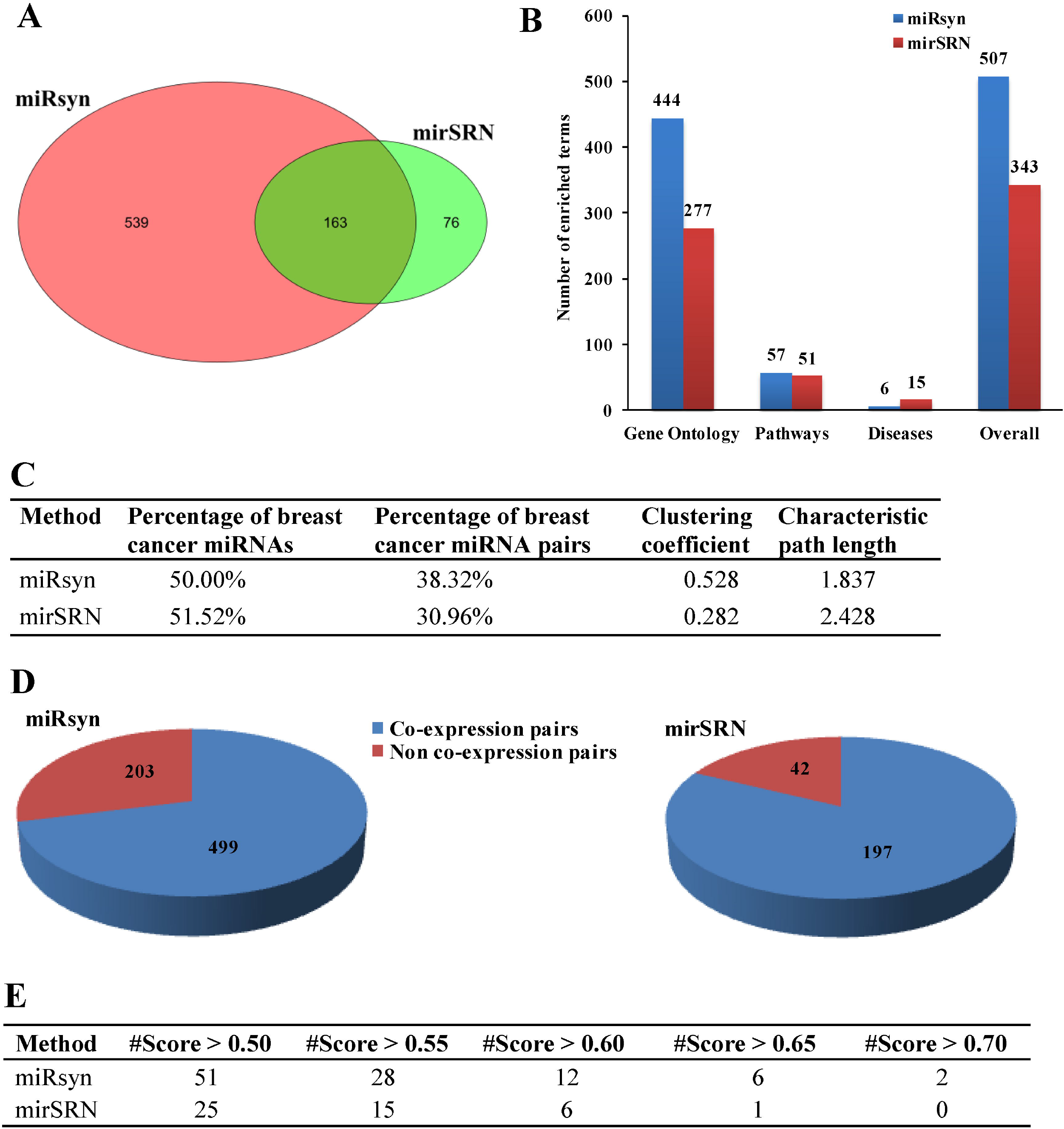
Comparison results between miRsyn and mirSRN. (A) The number of miRNA synergistic pairs. (B) The number of significantly enriched terms. (C) The percentage of breast cancer miRNAs and miRNA synergistic pairs, clustering coefficient and characteristic path length. (D) The number of co-expression and non co-expression miRNA synergistic pairs. (E) The overlap with putative miRNA synergistic pairs under different score cutoffs.

For the comparison in the percentage of breast cancer miRNAs and miRNA synergistic pairs (Figure 3C), the constructed miRNA synergistic network by mirSRN contains higher percentage of breast cancer miRNAs. However, the constructed miRNA synergistic network by mirSRN involves higher percentage of breast cancer miRNA synergistic pairs. Since the dense local neighborhoods and the small average characteristic path length can be exploited to predict miRNA synergism, Figure 3C implies that miRsyn is more suitable than mirSRN to identify miRNA synergism.

As shown in Figure 3D, most of synergistic miRNA-miRNA pairs identified by both miRsyn (~71.08%, 499 out of 702) and mirSRN (~82.43%, 197 out of 239) all show the same expression patterns. This comparison result indicates that the findings from miRsyn and mirSRN are consistent with each other. Although there is still no ground-truth for validating miRNA-miRNA synergistic pairs, we can use putative high-confidence miRNA-miRNA from the third-party database. In this work, we use the PmmR database [52] to compare the overlap with putative miRNA synergistic pairs between miRSyn and mirSRN. The score (the interval is [0 1]) in the PmmR database indicates the strength of each miRNA-miRNA synergistic pair, and a larger score denotes a stronger strength. Under different score cutoffs (range from 0.50 to 0.70 with a step of 0.05), the overlap of miRsyn is always larger than that of mirSRN (Figure 3E). This result indicates that several synergistic miRNA-miRNA pairs predicted by miRsyn (missed by mirSRN) still overlap with PmmR database. In sum, the above comparison results indicate miRsyn is more suitable than mirSRN in studying miRNA synergism.

## Conclusions

It is known that human complex diseases such as cancers are affected by multiple miRNAs rather than individual miRNA. Therefore, identifying miRNA synergism is important to understand the regulatory mechanisms of human complex diseases.

In this work, we have proposed a framework called miRsyn to identify miRNA synergism from both sequence and expression data. By using multiple-intervention causal inference, we simulated the causal effects of multiple miRNAs on target genes in the multiple-intervention experiments. To study miRNA synergism, we have conducted analysis at both network and module levels.

Topological analysis has shown that the constructed miRNA synergistic network is not scale-free, but small-world. The small-worldness may help the synergism of miRNAs to quickly adapt to a new biological environment caused by disturbances. In addition, most of synergistic miRNA-miRNA pairs show the same expression patterns, which allows for a rapid response to external disturbances.

We have also discovered that some miRNA-miRNA pairs at the sequence level are not working synergistically at the expression level. This result implies that it is necessary to study miRNA synergism from heterogeneous data sources. To further reveal the potential functions, we conducted functional enrichment analysis of synergistic miRNAs. The miRNA enrichment analysis results display that the identified miRNA synergistic network is directly or indirectly associated with the biological condition of breast cancer dataset. Moreover, by conducting breast cancer enrichment analysis, we have found that several miRNA synergistic modules are significantly enriched in breast cancer.

We compared our method miRsyn with mirSRN in different terms, including the number of synergistic miRNA pairs, the number of significantly enriched terms, the percentage of breast cancer miRNAs and miRNA synergistic pairs, clustering coefficient and characteristic path length, the number of co-expression and non co-expression miRNA synergistic pairs, and the overlap with putative miRNA synergistic pairs under different score cutoffs. The comparison results show that miRsyn (simulating multiple gene knock-down experiments) is more suitable than mirSRN (simulating single gene knock-down experiments) to identify miRNA synergism. For this current work, in order to have a fair comparison (i.e. using the same data types and similar inference method for estimating causal effects of miRNAs on mRNAs), we focus the comparison on one existing method mirSRN only. However, it is helpful to compare miRsyn to other different methods as well. In future, to further show the performance of miRsyn in studying miRNA synergism, we will conduct more comprehensive comparison.

Taken altogether, this work provides a novel framework to identify miRNA synergism that can be applied in variable biological fields. The presented results from the proposed method could provide insights to understand the synergistic roles of miRNAs in breast cancer. We believe that the presented method is applicable to the study of miRNA synergism associated with other human complex diseases.

## Abbreviations

miRNA: microRNA
nt: nucleotide
mRNA: Messenger RNA
PPI: Protein-protein interaction
GO: Gene Ontology
KEGG: Kyoto Encyclopedia of Genes and Genomes
TCGA: The cancer genome atlas
DAG: Directed Acyclic Graph
CPDAG: Completed Partially Directed Acyclic Graph
Joint-IDA: Joint Intervention calculus when the DAG is Absent
KNN: *k*-nearest neighbours

## Declarations

### Ethics approval and consent to participate

Not applicable.

### Consent for publication

Not applicable.

### Availability of data and materials

The datasets and source code in the current study are available at https://github.com/zhangjunpeng411/miRsyn.

## Authors’ contributions

JZ, NR and TDL conceived the idea of this work. VVHP, LL, TX, BT and JL refined the idea. JZ and VVHP designed and performed the experiments. TX, BT and JL participated in the design of the study and performed the statistical analysis. JZ, LL, JL, NR and TDL drafted the manuscript. All authors revised the manuscript. All authors read and approved the final manuscript.

## Acknowledgements

We would like to thank the reviewers for their valuable comments, which helped improve the work substantially.

## Additional files

### Additional file 1 –miRNA synergistic network

The miRNA synergistic network includes 702 miRNA-miRNA synergistic pairs, and 269 breast cancer related miRNA-miRNA pairs.

### Additional file 2 – miRNA enrichment analysis results of the synergistic miRNAs in the identified miRNA synergistic network

In total, we obtain 444 Gene Ontology terms, 57 Pathways terms and 6 Diseases terms related to the synergistic miRNAs, respectively.

### Additional file 3 –miRNA synergistic modules

The numbers of miRNA synergistic modules and breast cancer related miRNA synergistic modules are 361 and 72, respectively.

### Additional file 4 – Co-expression and non co-expression miRNA-miRNA synergistic pairs

The numbers of co-expression and non co-expression miRNA-miRNA pairs are 499 and 203, respectively.

### Additional file 5 – miRNA synergistic network at the sequence level

The number of miRNA-miRNA synergistic pairs is 1313 at the sequence level, and 611 synergistic miRNA-miRNA pairs of them are not working synergistically at the expression level.

### Additional file 6 – The comparison results of the mirSRN method

